# Utilizing comprehensive and mini-kinome panels to optimize the selectivity of quinoline inhibitors for cyclin G associated kinase (GAK)

**DOI:** 10.1101/471615

**Authors:** Christopher R. M. Asquith, Caleb Hopkins, Daniel K. Treiber, William J. Zuercher

## Abstract

We demonstrate an innovative approach utilising both fit-for-purpose kinase mini-panels and kinome-wide panels to progress discovery programs in the optimization of inhibitor potency and selectivity. We present a focused case study on development of a selective inhibitor of cyclin G associated kinase (GAK) using the quin(az)oline inhibitor chemotype. These results exemplify a versatile, efficient approach to drive kinome selectivity during inhibitor development programs.

## Introduction

A key challenge in developing kinase inhibitors that target the conserved ATP-binding site is inhibitor selectivity. The optimal screening approach would be to generate full kinome profiles in each lead series iteration and thus provide a complete picture of compound selectivity.^1^ Profiling every lead/compound throughout a medicinal chemistry program is a costly and time-consuming endeavour. There is a need to economise to have the optimal balance of time, cost, and information for progression. This in practice means testing a subset of kinases against a subset of compounds in the early stage. Indeed, there is increasing data suggesting that focused, rationally-designed kinome mini panels can provide useful, representative data for wider kinome selectivity to effectively support discovery efforts.^2-3^

We have now defined a compound progression sequence facilitated by the use of mini-panels at intermediate steps. At the outset, a broad evaluation of selectivity (KINOMEscan) would serve to de-risk and prioritize hit compound series. The use of fit-for-purpose KinaseProfiler mini-panels would allow for on target potency optimization while monitoring selectivity at each point. The optimization would conclude with a full kinome panel (KINOMEscan) for the final compound(s) and InCELL Pulse for cellular potency (**Figure 1**).

**Figure 1.**
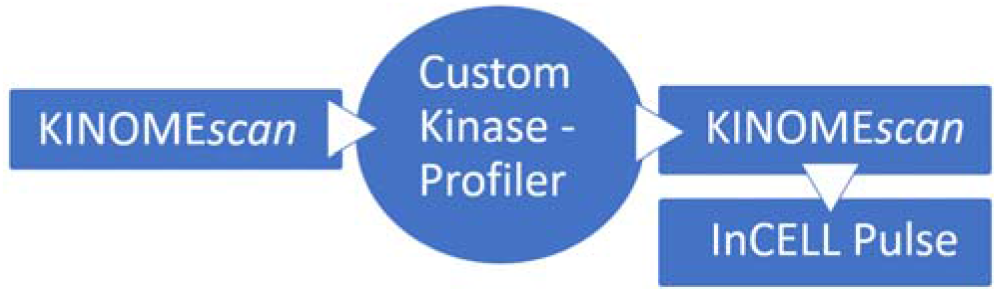
Proposed medicinal chemistry screening sequence for compound progression.

The quin(az)oline is a common kinase inhibitor chemotype. Several quinazoline-based clinical kinase inhibitors show cross reactivity with GAK, including the approved drugs gefitinib, erlotinib, and bosutinib (**Figure 2**).^4^ These drugs were designed as inhibitors of either epidermal growth factor receptor (EGFR) or SRC family kinases and show similar or higher affinity for several other kinases, making them ineffective tools for direct interrogation of GAK biology.^5^

*iso*thiazolo[5,4-*b*]pyridine (JMC 2015 - 12g) was reported as a selective GAK inhibitor.^6^ Although useful as a tool compound, JMC 2015 - 12g shows cross reactivity with other kinases, any of which may lead to confounding cell biology. The availability of a selective GAK chemical probe with a negative control would be a valuable asset for deconvoluting phenotypes associated with GAK.

Cyclin G associated kinase (GAK) is a 160 kDa serine/threonine kinase originally identified as a direct association partner of cyclin G and a member of the numb-associated kinase (NAK) family.^7^ In addition to its kinase domain, the GAK C-terminus has high homology with auxilin and tensin, with broad expression and localization to the Golgi complex, cytoplasm, and nucleus.^8-9^

GAK has been implicated in diverse biological processes, and genome-wide association studies have identified single nucleotide polymorphisms in the GAK gene associated with susceptibility to Parkinson’s disease.^10^ GAK is also involved in membrane trafficking and uncoating of clathrin-coated vesicles in the cytoplasm.^11,12^ GAK is also required for maintenance of centrosome maturation and progression through mitosis.^13^ GAK is over-expressed in osteosarcoma, and prostate cancer where GAK is implicated in progression to androgen independence.^14-16^ GAK inhibition has been associated with clinical toxicity due to pulmonary alveolar dysfunction based upon the phenotype of transgenic mice expressing catalytically inactive GAK, but this hypothesis has not yet been tested with a selective small molecule GAK inhibitor.^17^

**Figure 2.**
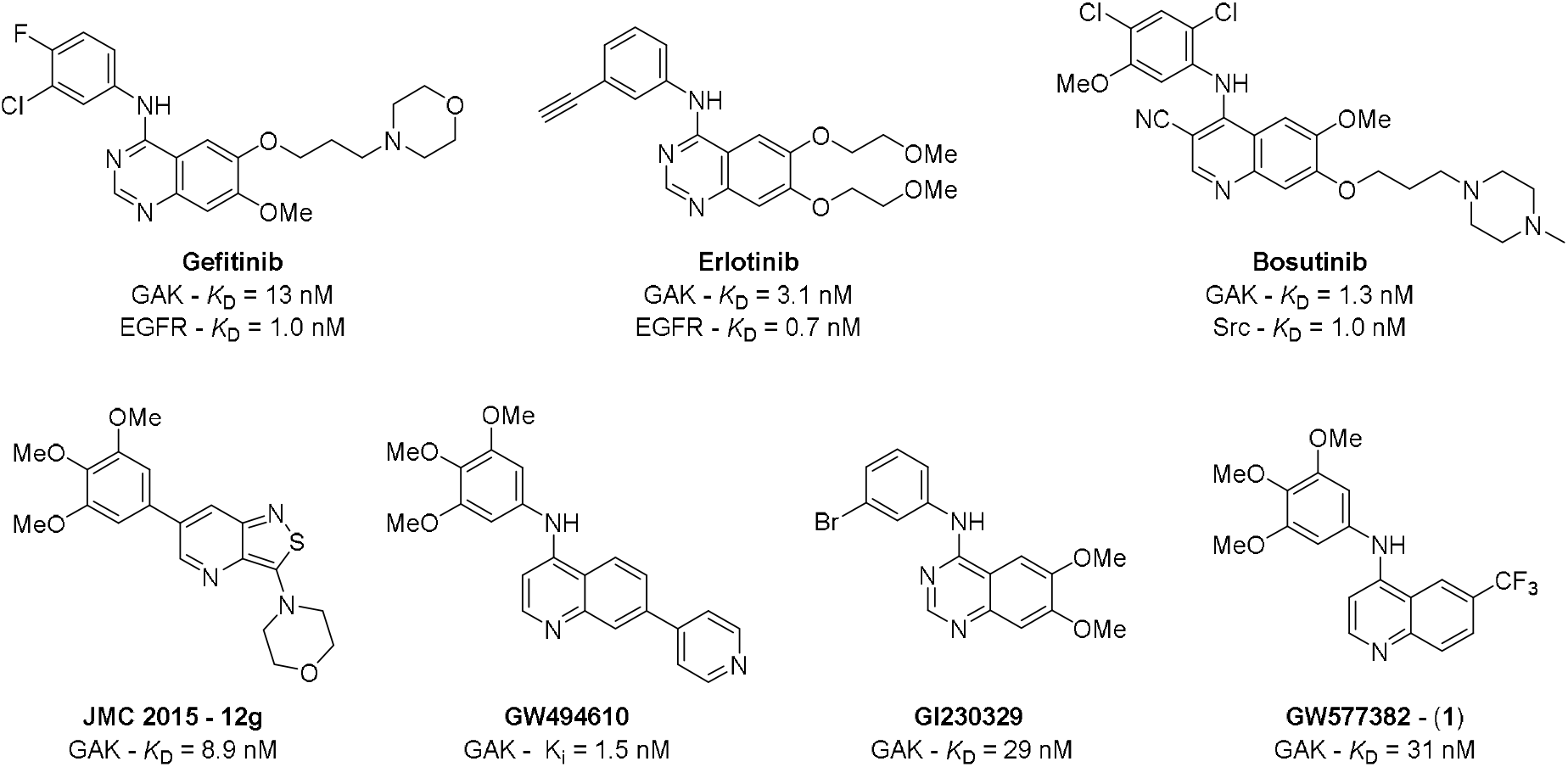
Previously described inhibitors of GAK.

We are focused on elucidating the biology of GAK and other members of the lesser-studied portion of the kinome.^18^ We have previously described the identification of a series of 4-anilinoquinoline inhibitors of GAK, exemplified by **1**.^19^ Quinoline **1** was profiled at 1 μM across an assay panel of over 400 wild-type human kinases (KINOMEscan). Subsequent *KD* determination for those kinases with > 60% binding at 1 μM identified three kinases (receptor-interacting protein kinase 2, RIPK2; AarF domain containing kinase 3, ADCK3; and nemo-like kinase, NLK) with *K_D_* values within 30-fold of that of GAK. We now elucidate the optimisation of **1** using fit for purpose KinaseProfiler mini-kinase selectivity panels.

## Results

We developed a series of GAK selective 4-anilinoquinolines and found that the trimethoxy aniline was the preferred substitution on the ‘head group’ of the molecule.^19,20^ This head group favourably balanced steric, electronic, and solvent effects. We now report a mini-panel kinome optimization of quinoline substitution using an activity-based KinaseProfiler assay format complementary to the on target GAK *K_D_* (**Table 1**). The unsubstituted quinoline (**2**) showed a >15-fold reduction in potency relative to the 6-CF_3_ analog **1**. The 6-position halogen-containing compounds **3**-**6** showed preference over those with halogens at the 7-position (**7**-**10**). The larger 6-*tert*-butyl (**11**) was 3-fold weaker than **1**, and 6-cyano (**12**) showing a doubling of potency. The 6-methylsulfone (**13**) was considerably weaker with more than 2 orders of magnitude loss in binding potency. The methoxy isomers (**14**-**16**) did not yield increased potency relative to the halogens, but **15** showed that the substitution pattern does affect GAK potency with a 3-5-fold improvement over mono-substituted analogs **14** and **16**. The 7-position cyano (**17**) and trifluoromethyl (**18**) showed no improvement over the corresponding 6-position counterparts **12** and **1** respectively.

We employed a custom panel of sixty-nine kinase activity assays (KinaseProfiler) to measure kinome-wide selectivity at 1 μM compound (**Table 1**, **Table S1 Figure 2**). This panel included all off-targets of **1** as well as a standard panel of diverse kinases. On the quinoline core, 7-position halogens (**8-10**) in addition to 7-trifluoromethyl (**17**) and 7-cyano (**18**) showed affinity for ABL. The same trend was observed with ALK2 with increased potency (80-90% inhibition) and to a lesser extent with ALK6. c-RAF was also potently inhibited across the series, with inhibition range of 70-90%. However, the monomethoxy compound **16** showed reduced activity by 20% on c-RAF showing that selectivity could be achieved.^21^ EGFR showed no significant activity at 1 μM across the series, despite extensive previous reports of EGFR activity on more decorated quin(az)oline structures.^4,5^

**Table 1.**
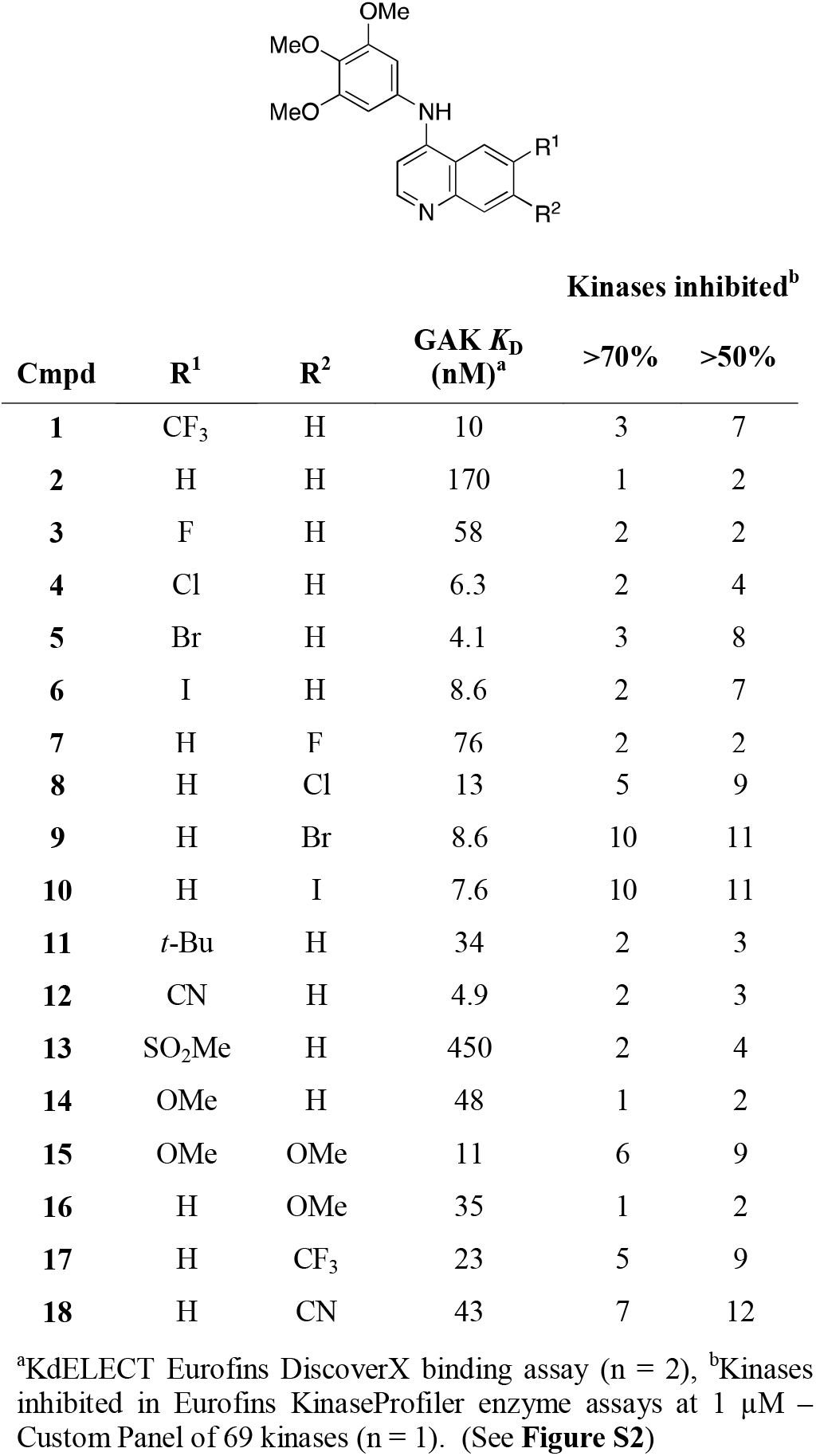
Affinity of 4-anilinoquinolines for GAK, and corresponding mini-kinome panel profile

| Cmpd | R<sup>1</sup> | R<sup>2</sup> | GAK $K_D$ (nM)<sup>a</sup> | Kinases inhibited<sup>b</sup> | |
| --- | --- | --- | --- | --- | --- |
|  |  |  |  | >70% | >50% |
| **1** | $\text{CF}_3$ | H | 10 | 3 | 7 |
| **2** | H | H | 170 | 1 | 2 |
| **3** | F | H | 58 | 2 | 2 |
| **4** | Cl | H | 6.3 | 2 | 4 |
| **5** | Br | H | 4.1 | 3 | 8 |
| **6** | I | H | 8.6 | 2 | 7 |
| **7** | H | F | 76 | 2 | 2 |
| 8 | H | Cl | 13 | 5 | 9 |
| 9 | H | Br | 8.6 | 10 | 11 |
| 10 | H | I | 7.6 | 10 | 11 |
| 11 | *t*-Bu | H | 34 | 2 | 3 |
| 12 | CN | H | 4.9 | 2 | 3 |
| 13 | SO<sub>2</sub>Me | H | 450 | 2 | 4 |
| 14 | OMe | H | 48 | 1 | 2 |
| 15 | OMe | OMe | 11 | 6 | 9 |
| 16 | H | OMe | 35 | 1 | 2 |
| 17 | H | CF<sub>3</sub> | 23 | 5 | 9 |
| 18 | H | CN | 43 | 7 | 12 |

<sup>a</sup>KdELECT Eurofins DiscoverX binding assay (n = 2), <sup>b</sup>Kinases inhibited in Eurofins KinaseProfiler enzyme assays at 1 μM – Custom Panel of 69 kinases (n = 1). (See **Figure S2**)

The ephrin receptor family showed broadly similar trends with EphA1, EphA8 and Fyn potently inhibited by 7-position halogenated compounds (**8**-**10**) in addition to 7-trifuoromethyl (**17**) and 7-cyano (**18**). EphA5 and EphB4 showed less sensitivity to changes in substitution but similar trends across the series. Interestingly the 6,7-disubstituted methoxy (**15**) was potent across the whole family as was the *meta*-methoxy 6-trifluoromethyl analog (**20**), suggesting that the structure activity relationship is not solely due to steric effects and is likely influenced by other contributions including electronic and solvation effects (**Figure 3**). Lyn and Lck followed the same trends as the ephrin receptor kinases, suggesting that substitution at the 7-position was increasing promiscuity more generally across the kinome.^19^ The last kinase to show sporadic inhibition was Txk with a profile consistent with the previous kinases albeit with reduced sensitivity.

**Figure 3.**
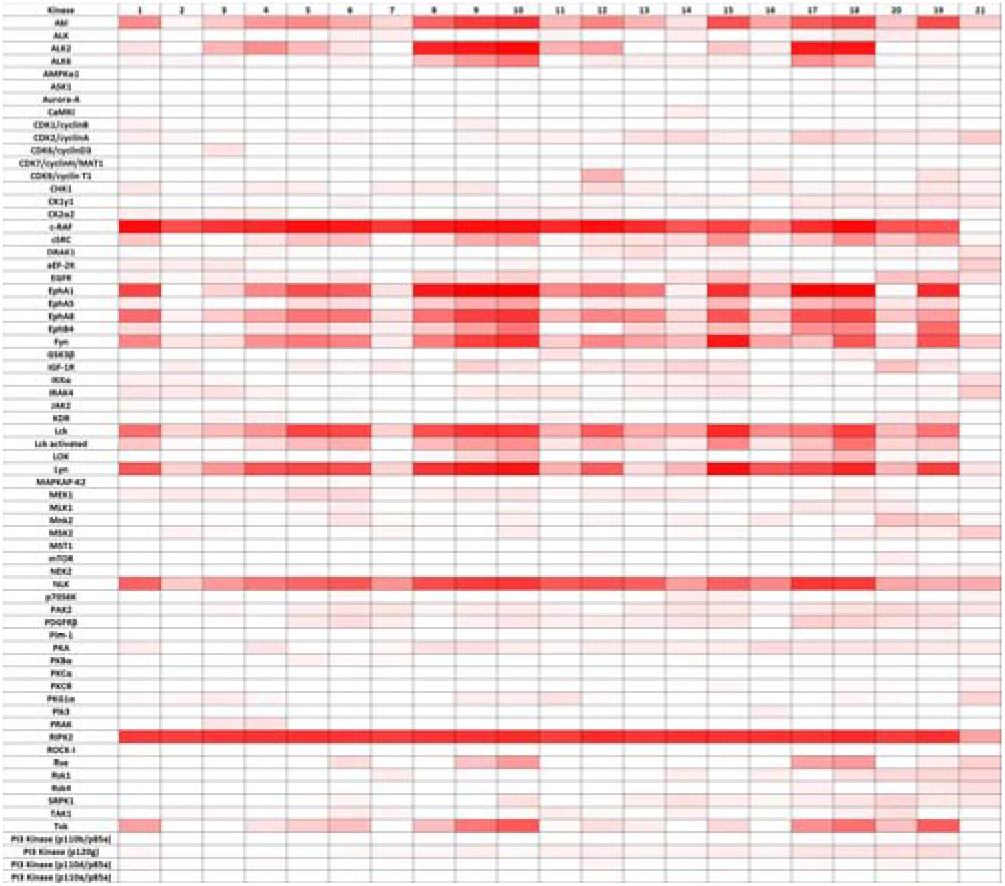
Eurofins KinaseProfiler customized panel heatmap on 69 kinases for 21 quin(az)olines (**See Table S1**)

The previously identified off-target nemo-like kinase (NLK) from the original hit compound **1** showed a range of profiles across the series with a pattern similar to the GAK activity. However, the RIPK2 activity was not sensitive to changes in substitution across the compounds tested. This observation held even with some of the more diverse analogs such as **19** and **20** but not with the negative controls **21** and **22** (**Figure 4**).

Utlising the results from the mini-panel panel structure actcity relationships, we chose compounds **5** and **15** for full-kinome follow-up to identify a candidate with the best GAK potency and overall selectivity profile on theKINOMEscan^®^ platform. (**Table 2**). We had initially screened quinoline **1** to reveal the kinome landscape with over 468 kinases screened, including 402 wild-type human kinases, 63 mutants, and 3 parasite kinases. Only GAK and three other off-targets (ADCK3, NLK & RIPK2) below 1 μM with 5 further kinases between 1-5 μM. The literature precedence around **15** and our previous report suggested this compound would be the next logical compound to screen across the kinome. However we found increased promiscuity across the kinome, particularly for ABL1, EphA6 and RIPK2 with the nearest off-target dropping from a 30-fold window to GAK in **1** to less than a 10-fold window for **15**. This was compounded by the wider range of thirteen off-targets compared to seven for **1** in the 0.1-5 μM range. We then screened compound **5** based on the mini-panel, which showed a narrow kinome profile and a corresponding potent GAK activity. We found the kinome selectivity profile had improved over **1** with increased potency on target.

**Figure 4.**
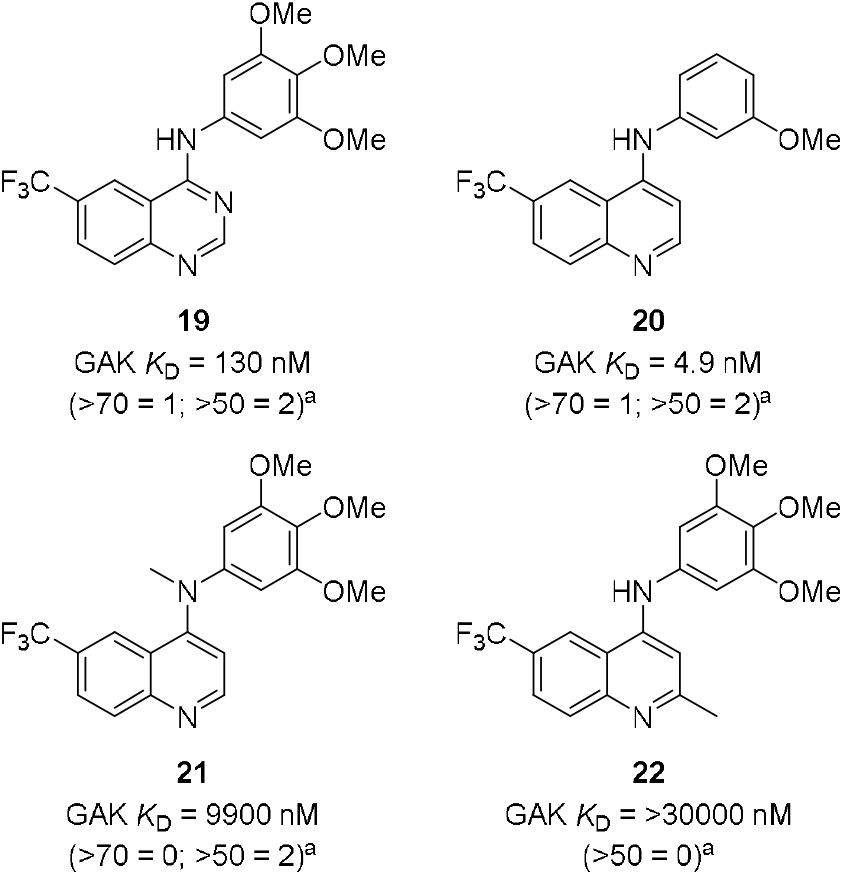
Analogs for further investigation of quinoline pharmacophore. ^a^Kinases inhibited Eurofins enzyme assay at 1 μM - Panel of 69 kinases (n = 1).

Analysis of the active site suggested that GAK Ala47 residue could be exploited to improve selectivity over RIPK2. In comparison to 6-cyano (**12**), the 6-bromo compound (**5**) was expected to have a reduced ability for a direct hydrogen bonding interaction between Ser25 of RIPK2 near the solvent exposed portion of the ATP-binding pocket.^22^ The corresponding negative control (**21**) only hit IRAK3 under 5 μM across the kinome. The kinome profiles of **1**, **15** and **5** show a progression to a compound that is both narrow spectrum and potent on target (**Figure 5**). The mini-panels enable an optimsation strategy driven by kinome selectively concurrent with kinase specific potency.

**Table 2.**
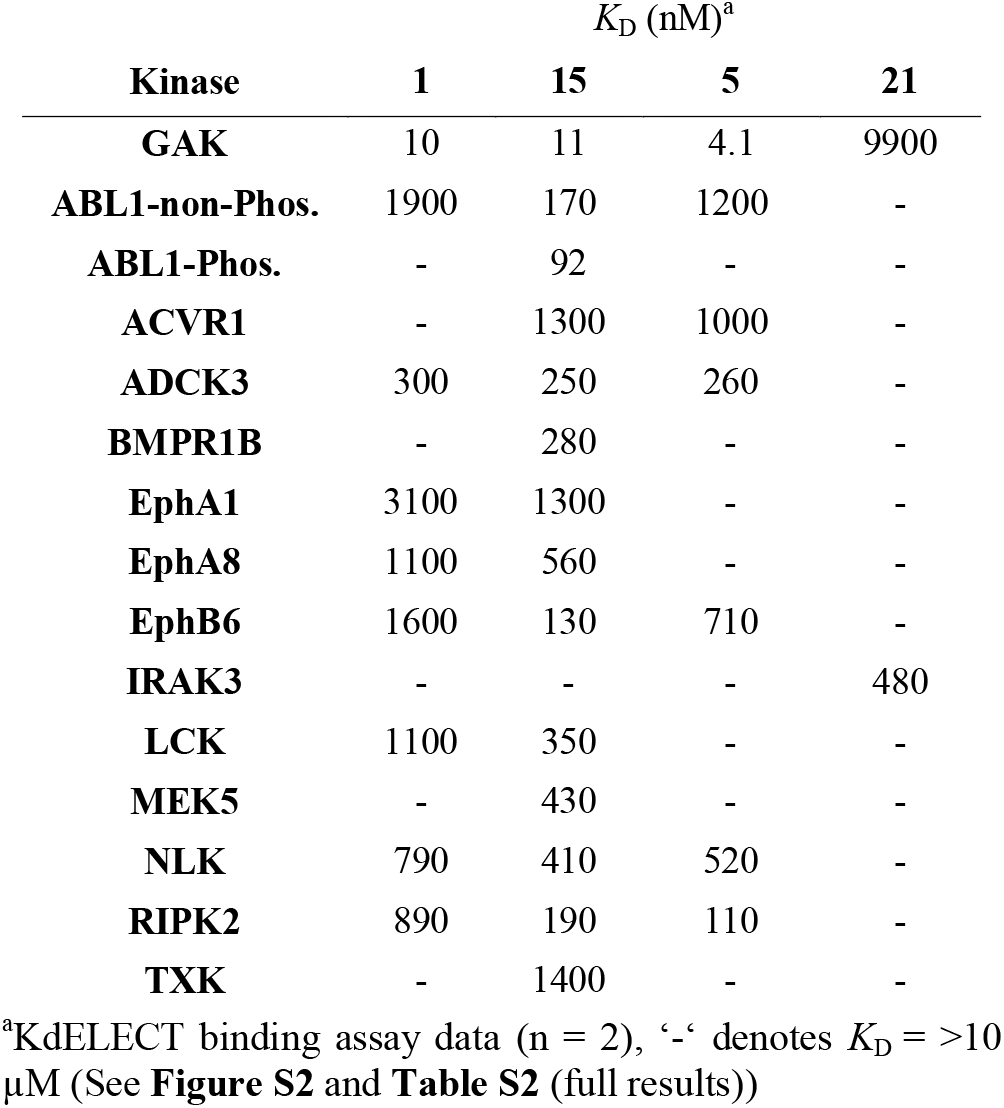
KINOME*scan* profile of 4 key compounds.

and direct readout of cellular compound permeability and target binding. We screened the most promising compounds in the InCELL Pulse, binding assay model cellular target engagement system.^23^ The compounds showed in cell activity against GAK, with good concordance with the *in vitro* binding (**Table 3**). The starting point **1** already had submicromolar activity (IC_50_ = 210 nM), with the final compound **5** showing similar potency. Compound **15** showed a > 4-fold improvement over **5** but with a mild increase in cross activity with additional kinases. Interestingly the *meta-methoxy* 6-trifluoromethyl analog **20** showed only a modest decrease in cellular potency, with a corresponding increase in ligand efficiency. The negative controls **21** and **22** performed as expected, with the strategically positioned additional methyl groups disrupting binding. The InCELL Pulse assay provided a robust read out of cellular potency (**Figure 6**).

**Table 3.** Cellular target engagement profile of seven key compounds

| Cmpd | GAK (nM)<sup>a</sup> | In-Cell (nM)<sup>a</sup> | Kinome<sup>b</sup> |
| --- | --- | --- | --- |
| 1 | 10 | 210 | 3 |
| 5 | 4.1 | 200 | 4 |
| 15 | 11 | 45 | 10 |
| 19 | 130 | 6700 | - |
| 20 | 4.9 | 380 | - |
| 21 | 9900 | >10000 | 1 |
| 22 | >30000 | >10000 | - |

<sup>a</sup>( $K_D$ ) KdELECT DiscoverX binding assay (n=2); <sup>b</sup>( $IC_{50}$ ) InCELL Pulse binding assay <sup>c</sup>scanMAX kinase hits under 1 μM (excluding GAK)

## Discussion

4-Anilinoquinolines and quinazolines are common structural templates for kinase inhibitors, including several approved medicines, with many more under clinical investigation.^4-5^ We had previously identified a series of 4-anilinoquinolines as narrow spectrum inhibitors of GAK and now show how we used mini-kinase panels to define several activity wells on different kinases within a focused array of compounds.

**Figure 5.**
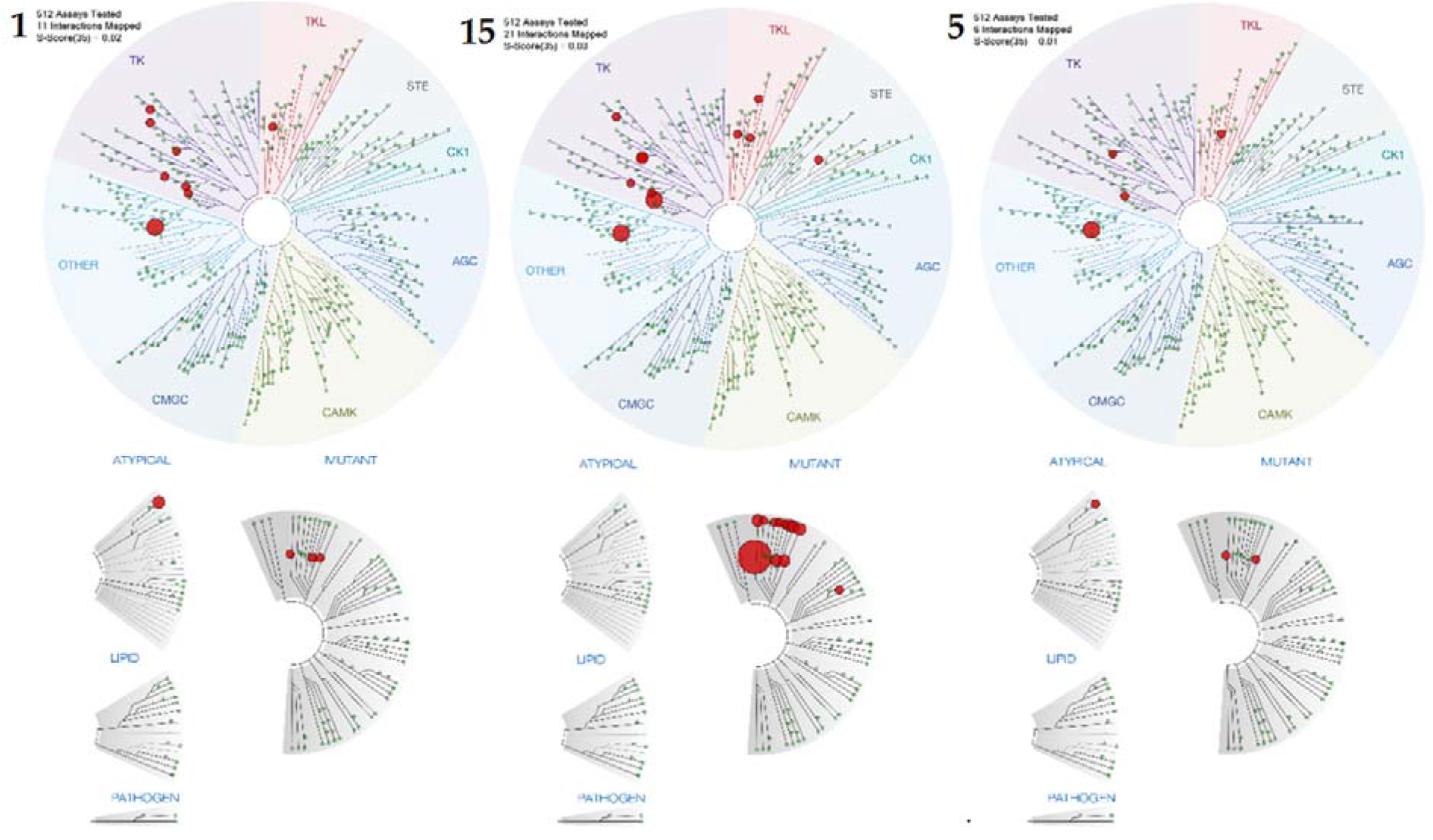
Kinome profiles (KINOME*scan*) of **1**, **15** and **5** visualization in TREE*spot*.

**Figure 6.**
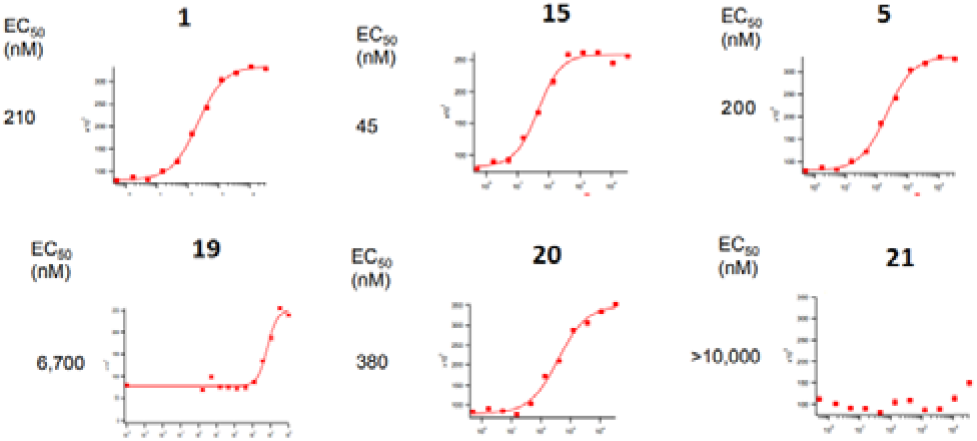
Eurofins-DiscoverX GAK InCELL Pulse IC_50_ curves for key compounds **1**, **15**, **5**, **19**, **20**, and **21** (see **Figure S3**).

The use of the full kinase panel, while ideal, can be both time and cost constrained during short medicinal chemistry cycles. The use of a focused mini-kinase panel is amenable to support early stage compound progression. This means promiscuous compounds are removed at an early phase of lead optimization, enabling compounds with a narrow kinome profile to progress for lead optimization. This approach also highlights the off-target structure activity relationships enabling them to be designed in or out.

We were able to see clear trends in the mini-panel results in tandem with the *scan*MAX results. We observed trackable off-targets in the mini-panel including ABL; c-RAF; NLK and RIPK2, but also more sporadic results within the ephrin receptor family (EphA1, A8) and other kinases including ALK2, Lck, Lyn and Txk. There were also clear trends within compound substitutions despite the solvent exposure of this region, with the 7-position showing more promiscuity than the 6-position except for **15**. It was also interesting to see several one-off weaker hits. These included the 7-cyano (**13**) on CDK9 but no other CDKs which is consistent with senexin A and B as narrow spectrum CDK inhibitors.^24^ The switch from the quinoline (**1**) to quinazoline (**19**) saw a slight increase in EGFR activity; further evidence of a complex interplay of structure activity relationships in kinase inhibitor design.

This work highlights the subtleties in kinase inhibitor design where small changes can have a large impact on the kinome selectivity profile. This study illustrates that a carefully crafted, fit for purpose combination of full- and mini kinome panels can enable efficient discovery of potent and selective inhibitors.

## Experimental

### Chemistry: General procedure for the synthesis of 4-anilinoquinolines

6-Bromo-4-chloroquinoline (1.0 eq.), 3,4,5-trimethoxyaniline (1.1 eq.), and ^i^Pr_2_NEt (2.5 eq.) were suspended in ethanol (30 mL) and refluxed for 18 h. The crude mixture was purified by flash chromatography using EtOAc:hexane followed by 1-5 % methanol in EtOAc. After solvent removal under reduced pressure, the product was obtained as a free following solid. Compounds **1**-**7** and **11**-**22** were synthesised as previously reported.^19,20^

### 7-chloro-*N*-(3,4,5-trimethoxyphenyl)quinolin-4-amine

(**8**) yellow solid (185 mg, 0.538 mmol, 71 %) MP 308-310 °C; ^1^H NMR (400 MHz, DMSO-d_6_) δ 11.21 (s, 1H), 8.92 (d, J = 9.1 Hz, 1H), 8.48 (d, J = 7.0 Hz, 1H), 8.22 (d, J = 2.1 Hz, 1H), 7.84 (dd, J = 9.1, 2.1 Hz, 1H), 6.87 (d, J = 7.0 Hz, 1H), 6.83 (s, 2H), 3.80 (s, 6H), 3.72 (s, 3H). ^13^C NMR (101 MHz, DMSO-d_6_)δ 155.0, 153.6 (s, 2C), 143.2, 139.1, 138.2, 136.6, 132.6, 127.2, 126.2, 119.2, 115.8, 103.2 (s, 2C), 100.8, 60.2, 56.2 (s, 2C). HRMS m/z [M+H]^+^ calcd for C_18_H_18_N_2_O3Cl: 345.1006, found 345.0996, LC t_R_ = 3.46 min, >98% Purity.

### 7-bromo-*N*-(3,4,5-trimethoxyphenyl)quinolin-4-amine

(**9**) yellow solid (189 mg, 0.483 mmol, 78 %) Decompose >250 ^o^C; ^1^H NMR (400 MHz, DMSO-d_6_) δ 11.21 (s, 1H), 8.83 (d, J = 9.1 Hz, 1H), 8.47 (d, J = 7.0 Hz, 1H), 8.37 (d, J = 2.0 Hz, 1H), 7.95 (dd, J = 9.1, 2.0 Hz, 1H), 6.87 (d, J = 7.0 Hz, 1H), 6.83 (s, 2H), 3.80 (s, 6H), 3.72 (s, 3H). ^13^C NMR (101 MHz, DMSO-d_6_)δ 155.1, 153.6 (s, 2C), 143.0, 139.1, 136.6, 132.6, 129.8, 127.2, 126.0, 122.3, 116.0, 103.2 (s, 2C), 100.8, 60.2, 56.2 (s, 2C). HRMS m/z [M+H]^+^ calcd for C_18_H_18_N_2_O_3_Br: 389.0501, found 389.0494, LC t_R_ = 3.53 min, >98% Purity.

### 7-iodo-*N*-(3,4,5-trimethoxyphenyl)quinolin-4-amine

(**10**) yellow solid (176 mg, 0.404 mmol, 78 %) MP 308-310 ^o^C; ^1^H NMR (400 MHz, DMSO-d_6_) δ 11.12 (s, 1H), 8.60 (d, J = 9.0 Hz, 1H), 8.53 (d, J = 1.7 Hz, 1H), 8.44 (d, J = 7.0 Hz, 1H), 8.08 (dd, J = 8.9, 1.7 Hz, 1H), 6.87 (d, J = 7.0 Hz, 1H), 6.82 (s, 2H), 3.80 (s, 6H), 3.72 (s, 3H). ^13^C NMR (101 MHz, DMSO-d_6_)δ 155.2, 153.6 (s, 2C), 142.7, 138.9, 136.6, 135.2, 132.63, 128.4, 125.2, 116.2, 103.2 (s, 2C), 102.0, 100.6, 60.2, 56.2 (s, 2C). HRMS m/z [M+H]^+^ calcd for C_18_H_1_N_2_O3I: 437.0362, found 437.0346, LC t_R_ = 3.70 min, >98% Purity.

### Kinase binding assays, Kinase enzyme assays KINOME*s-can* assays and *scan*MAX assays

preformed as previously described. All assays were provided by Eurofins-DiscoverX (see SI).^1,23^

### InCELL Pulse

cellular target engagement assays were designed and performed according to manufacturer’s instructions (Eurofins-DiscoverX, Fremont, CA. Cat. No. 94-40075).^24^

### Supporting Information

The Supporting Information is available free of charge on the ACS Publications website at DOI: XXXXX. Experimental protocols and supporting figures and tables (PDF)

## Author Contributions

The manuscript was written through contributions of all authors. All authors approved of the final version of the manuscript.

## Notes

The authors declare no competing financial interests.

## ACKNOWLEDGMENT

The SGC is a registered charity (number 1097737) that receives funds from AbbVie, Bayer Pharma AG, Boehringer Ingelheim, Canada Foundation for Innovation, Eshelman Institute for Innovation, Genome Canada, Innovative Medicines Initiative (EU/EFPIA) [ULTRA-DD grant no. 115766], Janssen, Merck KGaA Darmstadt Germany, MSD, Novartis Pharma AG, Ontario Ministry of Economic Development and Innovation, Pfizer, São Paulo Research Foundation-FAPESP, Takeda, and Wellcome [106169/ZZ14/Z]. We also thank Dr. Brandie Ehrmann for LC-MS/HRMS support provided by the Mass Spectrometry Core Laboratory at the University of North Carolina at Chapel Hill and Alex Lun for technical support.

## Table of Contents Graphic

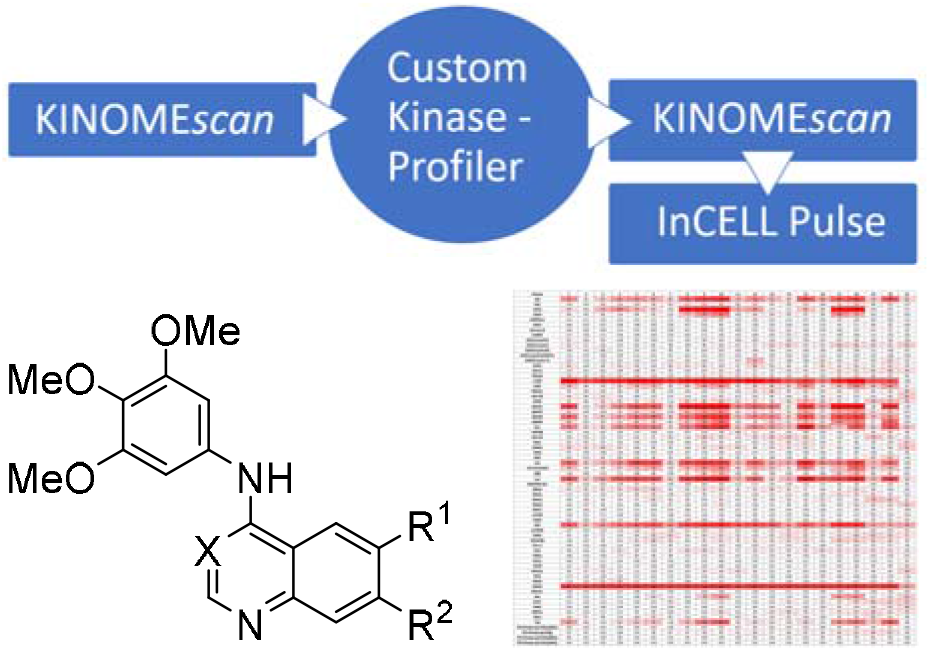

